# Expanding the bioluminescence reporter toolkit for plant chronobiology with NanoLUC

**DOI:** 10.1101/439844

**Authors:** Uriel Urquiza-García, Andrew J. Millar

## Abstract

Bioluminescence has been an important tool for gathering circadian data with the main reporter gene exploited firefly luciferase (LUC). In some circumstances the rapid inactivation of LUC could be disadvantageous, e.g. reporting total protein levels through reporter translational fusions. In the latter scenario the commercially available Nano luciferase (NanoLUC) might offer and advantage, however no data in plant has been provided so far. We tested NanoLUC under different research scenarios were LUC has been used, for example enzyme purification, expression in transient plant systems and in stable transgenic lines. We show that NanoLUC is active in these experimental scenarios. We also created a set of NanoLUC variants for example MBP-NanoLUC-3xFlag-10xHis version of NanoLUC can be easily purified and stable for several days (half-life 37.2 at 4°C) and can be used for generating calibration curves for quantifying protein as the signal is linear over a large dynamic range. In addition, we show that NanoLUC can report in-planta protein levels on circadian time scale thanks to the stability of furimazine. Therefore, opening the possibility of using NanoLUC for reporting protein dynamics in seedlings. With this new technology, we explored the dynamics of protein BROTHER OF LUX ARRHYTHMO (BOA), which has been suggested in mathematical models to be a rhythmic protein from RNA data. Using an automatic plate-reader, we were able to track BOAp:BOA-NL for an extended period of time by just adding a mix of furimazine with Triton X-100, as it is done with LUC-reporter lines. In our experimental context BOA protein does not present strong oscillatory dynamics similar to what has been reported for Phytocrhome B for which rhythmic accumulation of transcript can be observed while total protein levels remain constant under diurnal conditions. Our results support the use NanoLUC for studying the dynamics of plant proteins for extended period of time under different circumstances.

## Introduction

The use of firefly luciferase (*LUC*) as a reporter gene for plant chronobiology has been crucial for gaining insight in the genetic bases of circadian rhythmicity in plants and other model organisms. This was achieved first in Arabidopsis by fusing to the promoter of *CHLOROPHILE A/B BIDNING PROTEIN 2 (CAB2)*. The resulting fusions presents oscillations of light emission when luciferin is supplemented externally and can be tracked for several days in different environmental conditions. This powerful methodology allowed the isolation of the first circadian mutants in *Arabidopsis, TIMING OF CAB EXPRESSION (TOCs*) (Millar, Short, Hiratsuka, et al. 1992; Millar et al. 1995). The power of this assay is linked to the very short half-life of LUC, providing a faithful transcriptional readout of transcriptional activity. Additionally, modified versions of LUC have been exploited for studying even faster dynamics, such as transcriptional bursts in other systems (Suter et al. 2011). The reporter has been used for gathering timing data and waveform characteristics.

Quantifications of LUC signal levels relative to total protein content have been performed of for CIRCADIAN CLOCK ASSOCIATED 1 (CCA1) and TIMING OF CAB EXPRESSION 1 (TOC1) in *Ostreococcus tauri*. (Corellou et al. 2009). However, thermal inactivation of LUC could result in underestimation of total protein levels even if cofactors are present in optimal conditions. It is important therefore that the reporter presents high-stability and its degradation is completely determined by the stability of the protein to be reporter. A more stable luciferase would provide, in principle, a better system for quantitatively studying protein levels both *in-vitro* and *in-planta*. Nano Luciferase (NanoLUC, NL) from Promega might provide a better option for this scenario (Hall et al. 2012). The reported data for NanoLUC suggest that it might provide a new tool set for studying protein dynamics on circadian time scale (5-10 days). However, this has not been reported elsewhere, therefore we tested the compatibility of NanoLUC in a lab environment that has exploited the use of LUC for gaining powerful insight into the genetics of circadian rhythmicity in *Arabidopsis*.

NanoLUC is a small protein of 19.2 kDa from the deep-sea shrimp *Oplophorus gracilirostris*. It was engineered for high stability (t_1/2_=11.5 days at 37°C) and codon optimised for expression in mammalian cells (Hall et al. 2012). Conveniently, the expected Codon Adaptation Index for expression in *Arabidopsis* is remarkably good eCAI = 0.737 (p<0.05), as calculated by E-CAI (Puigbò et al. 2008), on a scale where 1 is a sequence with the most common codons in the organism. The enzyme only requires O_2_ and substrate. The NanoLUC substrate is Furimazine, a Coelentarzine analogue, that presents a higher stability than luciferin. Also, the enzyme shows stable light emission around 22°C and stable activity in the physiological pH for Arabidopsis (7.1 in nucleus, 7.2 in cytosol (Shen et al. 2013)). This contrasts with LUC activity which is significantly more sensitive to changes in temperature and pH. LUC suffers product inhibition, resulting in a ‘dark’ (inactive) pool of enzyme after it is exposed to luciferin (Millar, Short, Chua, et al. 1992). It also requires two cofactors ATP and Mg^++^, which might fluctuate in a circadian manner (Feeney et al. 2016); whereas NanoLUC has not been shown to present this issue. If the activity of LUC is determined in plant extracts excess of cofactors can be supplied for eliminating any rhythmic behaviour in this species. These drawbacks in LUC compromise the determination of protein levels *in-vivo*, which will be an important addition to the bioluminescent tool-set. Finally, NanoLUC has been reported to generate light signal three orders of magnitude brighter than LUC (Hall et al. 2012). In principle this would allow for lower number of molecules to be detectable, however this has not been tested for *in-vivo* activity in plant systems. We decided to track the dynamics *BROTHER OF LUX ARRHYTHMO (BOA)*, also known as *NOX*. Chip-PCR data using ELF3 at the *PRR9* promoter in different mutant backgrounds suggest that BOA promotes binding of the Evening Complex by working together with *LUX ARRHYTHMO (LUX*) (Chow et al. 2012). In the clock mathematical model of Arabidopsis thaliana F2014 the authors introduced BOA and the recovered dynamics after fitting the model parameters predict the BOA protein levels are rhythmic (Fogelmark & Troein 2014). However, little data has been reported in the literature with only one western blot using an antibody against the native protein reported (Dai et al. 2011). In order to extend this work we created a collection of Col-0 CCA1p:LUC BOAp:BOA-NanoLUC and Col-0 BOAp:BOA-LUC. We found a discrepancy between the amplitude of BOA-NanoLUC and BOA-LUC and mathematical modelling suggests a differential stability between two reporters, with NanoLUC being more stable. Our current data suggest that BOA does not cycle with strong amplitude at the protein level contrary for the data reported for its transcript. This type of result has been observed for *PHYTOCHROME B (PhyB)* where a Phybp:LUC and PhyBp:PhyB-LUC present circadian oscillation. However, total PhyB protein levels do not present oscillatory behaviour judged by western blot experiments (Bognár et al. 1999). Through simple mathematical exercise we explore the source of this discrepancy between BOA constructs. Our results suggest that this discrepancy could be explained by differences in the stability of LUC and NanoLUC reporters. However, there may be other explanations which are considered in more detail within the discussion.

## Methods

### Construction of plasmids and bacterial work using NanoLUC

Plasmids pNL1.1 and pNL1.2 with NanoLUC sequence were provided by Promega corporation and propagated in DH5alpha (Invitrogen). NanoLUC sequence was amplified adding a NcoI/BamHI sites and cloned into pET28a(+) using the same sites. The NanoLUC sequence was again amplified and cloned in a pUC19 carrying a 3× Flag 10×His peptide forming a translational fusion NL-3xFLAG-10xHis (NL3F10H). This sequence was then amplified and cloned using Gibson assembly into pET28a(+) resulting in pET28aNL3F10H. The addition of then a N-terminal Maltose Binding Protein 6xHis-NanoLUC-3F10H was achieved by amplifying the MBP-6xHis from pMJ806 which was originally carrying a CAS9 protein and fused to NL3F10H and pET28a(+) resulting in pET28a::MBP-NL310H (Jinek et al. 2012). Each intermediate amplified step was sequence-verified after cloning by Sanger sequencing (Genepool, Edinburgh Genomics).

### NanoLUC purification protocol for calibration curves

1. Three days before creating a NanoLUC calibration curve, transform the vector pET28a(+)∷MBP-NanoLUC-3F10H using chemically competent cells BL21 DE3 Rossetta2 pLysS, selecting on LB with 50 μg/ml Kanamycin (Kan^50^) and 34 μg/ml Chloramphenicol (Cm^34^). Incubate the plate over-night at 37°C.
2. Pick a single colony and inoculate a 5 ml of LB with 50 μg/ml Kan, 34 μg/ml Cm and incubate at 37°C, 200 rpms overnight.
3. Early next day, inculcate 100 mL of LB Kan^50^ Cm^34^ in a 0.5 L Erlenmeyer to a final O.D._600nm_ of 0.01. Incubate at 37°C, 200 rpms following the growth until the O.D. has reached 0.5.
4. When the culture reached this point add IPTG to a final concentration of 1 mM and allow induction to proceed at 30°C for 8 hours.
5. Transfer the flask in ice-cold water for 10 min and then split the culture into two 50 mL polypropylene conical tubes. Harvest the cells by centrifugation at 4,000 r.f.c. 4°C for 15 min. and discard the supernatant.
6. Resuspend the cell in 10 ml ice-cold Lysis buffer (50mM NaH_2_PO_4_, 300mM NaCl, 10mM Imidazole, pH 8.0 NaOH adjusted).
7. Then sonicate the cells with by sonication conditions 1 min with 10s on and 10 off and 50% amplitude in ice.
8. Pass the crude lysate several time a 25G syringe needle to reduce lysate viscosity generated by genomic DNA.
9. Centrifuge 2 ml of lysate at 20,000 rfcs for 20 min at 4°C, which will eliminate unbroken cells and some large debris.
10. Transfer 1ml of supernatant to a 2 μl tube (safelock, Eppendorf), and mix with 250 μl of Ni-NTA agarose (Qiagen), incubated at 4°C for one hour with gentle agitation.
11. Recover the Ni-NTA beads by centrifugation at 20,000 rfc for 1 min transfer the supernatant to a clean 1.5 polypropylene tube for later analysis by SDS-PAGE.
12. Wash the Ni-NTA agarose beads three times with Washing Buffer (50 mM NaH_2_PO_4_, 300 mM NaCl, 20mM Imidazole, pH 8.0 NaOH adjusted), centrifuged 20,000 rfcs each time.
13. Elute the MBP-NL-3F10 with Elution Buffer (50 mM NaH_2_PO_4_, 300 mM NaCl, 250 mM Imidazole, pH 8.0 NaOH adjusted).
14. Dialyse the elution faction using a 10,000 MW cut-off membrane overnight at 4 °C in BI buffer (50 mM NaH_2_PO_4_, 300mM NaCl, 20 mM pH 8.0 NaOH adjusted).
15. Determine NanoLUC activity using a 1×10^−3^ dilution in BI buffer by taking 20 μl of diluted enzyme mixing it with 80 μl of BI buffer and 100 μl of 1:50 furimazine:NanoGlow assay buffer as a reference of starting activity.

### Generation of Gateway Binary Vectors for C-term NanoLUC fusions

Agrobacterium vectors pGWB604, pGWB605, pGWB601, pGWB701 and series carrying LUC+ for C-terminal translational fusions pGWB634, pGWB635, pGWB636, pGWB737 (Nakamura et al. 2010; Nakagawa et al. 2007) and propagated in E. coli ccdB resistant strain (New England Biolabs). Using the same design as pGWB63x we created an equivalent NL3F10H version instead of LUC+. pGWB60xNL3F10H. This was achieved by amplifying a sub-fragment of the gateway cassette and fusing it with NL3F10H and pGWB601 digested with NcoI/SacI by Gibson assembly. The CaMV35S promoter was amplified from pCAMBIA1305.1 adding attB1 and attB2 sites recombined into pDONR221 sequence verified and then introduced into pGWB601NL3F10H and pGWB635 (LUC+) by a Gateway LR reaction (Invitrogen). The genomic sequence coding for of BOA from −1000 to + 1053 (BOAp:BOA) without stop codon was amplified and cloned into the NL3F10H and LUC vectors in with a similar method as 35S promoter.

### Plant transformation and selection of homozygous lines

Col-0 or Col-0 CCA1p:LUC were stratified for two days at 4°C then transferred to 16L:8D at ~100 μmol/cm^2^s warm fluorescent light bulbs. After one month in these conditions, plants were decapitated from promoting branching and dipped three times with one-week interval between dipping with *Agrobacterium ABI* strain carrying 35S and BOA constructs described above. T0 seed was collected and primary transformants selected using BASTA resistance. The collection was then screened for segregation 3:1 on ROBUST media (1% Agar (Sigma), 1/2 MS, pH 5.8 NaOH adjusted) supplemented with 10 μg/ml Bilaphos (Sigma). Lines presenting single insertions were taken to homozygocity by selecting for 100% BASTA resistance on soil in T2 generation.

#### Plant material and Growth conditions for luciferase in-plants assays

For imaging experiments with LUC and NanoLUC seeds were surface sterilised and individual seeds placed in a flat white 96 well plate Lumitrac (Grenier) containing 150 μl of ROBUST media (1% Agar (Sigma), 1/2 MS, pH 5.8 NaOH adjusted). The plate containing seeds was stratified for 48 hours at 4 °C, then transferred to entrainment conditions (12L:12D 21°C ~100 μmol/cm^2^s fluorescent white cool light). After one week 50 μl of either 1:50 Furimazine:0.01% Triton X-100 (NanoLUC) or 5 mM Luciferin in 0.01% Triton X-100 (LUC) was added to each well. Plates were transferred to a 12L:12D photoperiod in light conditions 50 μmol/cm^2^s (monochromatic red and blue light) at 21°C under a 12L:12D photoperiod for three says and the transferred to constant conditions. Light emission was measured during this period every 30 min with an integration time of 1.5s using a Tristar plate reader (Berthold). Period analysis was performed in www.Biodare2.ed.ac.uk using FFT-NLS {Zielinski:2014dy}.

## Results

### Activity and Purification of NanoLUC variants

In order to verify the activity of the NL we cloned the sequence provided by Promega from pNL1.1 into pET28a(+) and performed an induction experiment in *E. coli* BL21 Rosetta2 with 1mM IPTG. After 6 hours of induction at 30 °C, cells were harvested by centrifugation, re-suspended in 1 ml of Phosphate Buffered Saline (PBS) and 1μl of Furimazine (NL substrate, stock concentration). Cells carrying the NanoLUC presented bioluminescence, while cells empty pET28a(+) did not (Figure 1.A). We created two NanoLUC derivatives. A synthetic 3xFlag 10xHis tag was added on the C-terminus of NanoLUC in pET28 resulting in NL3F10H. Addition of the 10xHis allows purification by Ion Metal Affinity Purification (IMAC) using a Ni-NTA matrix as a stationary phase. Also, an MBP-NL-3F10H (Maltose Binding Protein, MBP) was created to test the effect of fusing proteins to the N-terminus of NL. After the MBP, a 6xHis tag was also added, so this NL presents 16xHis. The resulting constructs were introduced into *E. coli* BL21 Rosetta 2 pLysS for expression and purification. The three versions NL, NL3F10H and MBP-NL3F10H presented high expression levels at 30 °C and remained soluble after a centrifugation for 20 min a 20,000xg (Figure 1.B, lysate lane). We performed an IMAC purification using Ni-NTA agarose beads (Qiagen) with the manufacturers standard protocol, resulting in successful isolation. As expected the NL original variant does not bind to Ni-NTA beads and can be removed after two washes with Washing Buffer (WB) (Figure 1.A). The NL3F10H version did bind to the Ni-NTA beads and eluted with high Imidazole concentrations present in the Elution Buffer (EB) (Figure 1.B). This shows 3F10H tag allows purification of NL resulting in a homogenous preparation that can be used for generating standard curves. As expected for MBP-NL3F10H a very significant band shift is observed and a homogenous purification can be observed in the PAGE gels (Figure 1.C). This shows that purification of NL can be performed in a straight forward way for several uses including generation of calibration curves for inferring NanoLUC concentration in plant extracts.

**Figure 1.**
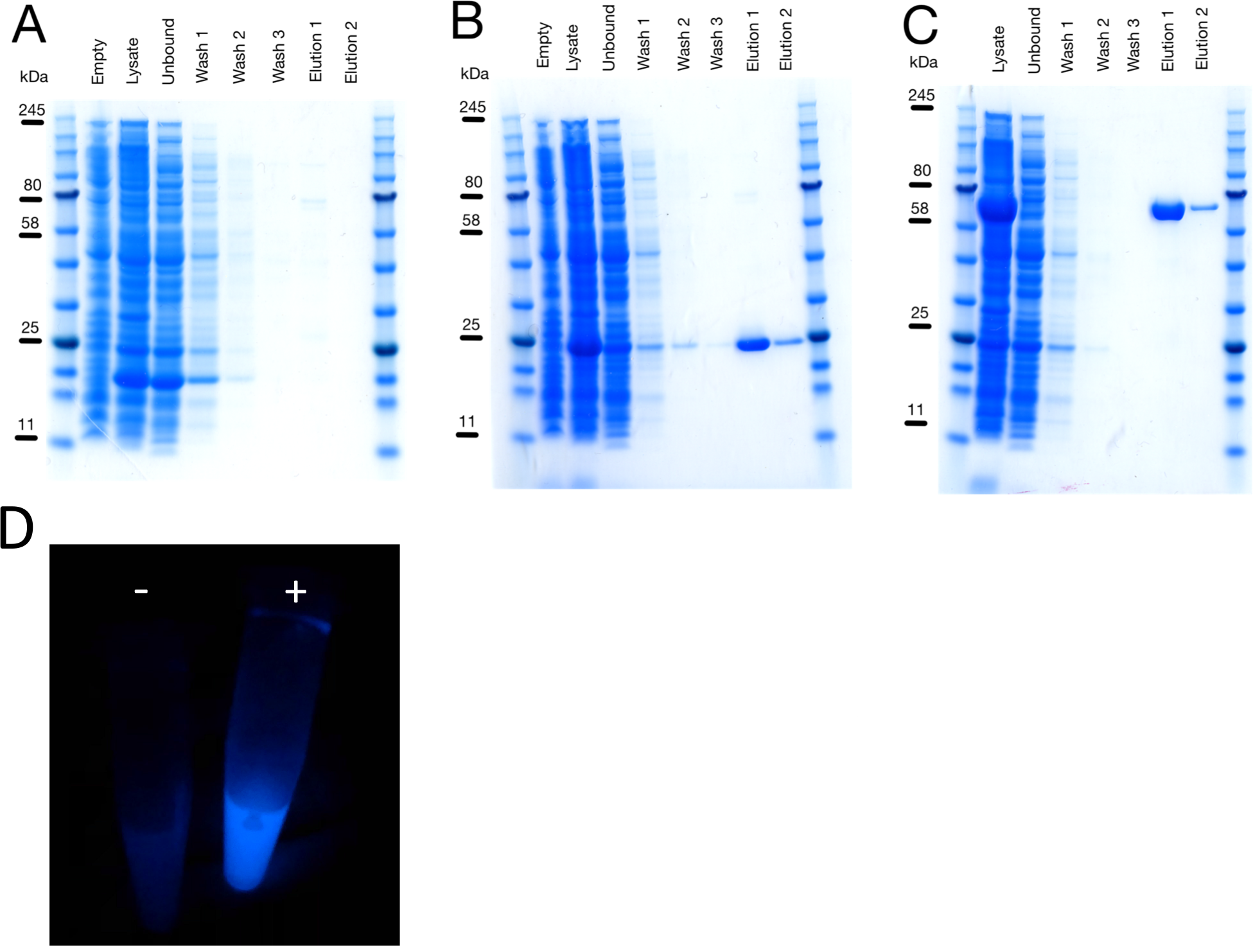
Recombinant expression of NanoLUC variants. A) NanoLUC provided by Promega expressed in Rosetta 2 BL21 strain induced with 1mM IPTG at 30°C. B) NanoLUC 3xFLAG 10xHis. C) MPB-NL3F10H 10 μl of each fraction were loaded into a NuPAGE 4-12% Bis-Tris and run at 200V for 40 min. All the variants were produced as in A. Purification was performed as described in the in methods. D) Tubes containing cells induced with IPTG (+) and without being induced (−) NanoLUC emits blue light, to exemplify the brightness of NanoLUC the photograph was taken with standard mobile phone camera.

Then we compared the activity of NL3F10 to MBP-NL3F10H in order to test detrimental effects on activity by adding this N-terminal tag. This comparison is important because we will be using NL3F10H as C-terminal tag for reporting the levels of transcription factors in stable transgenic plants. We purified NL3F10H and MPB-NL3F10H variants as before, however a dialysis step was added in order to remove the excess of Imidazole used for eluting the protein from the Ni-NTA beads. The dialysis was performed at 4°C over-night. This step was introduced because Imidazole can interfere with the Bradford protein assay, resulting in poor protein quantifications. The elution fractions for the two variants were protein quantified using a linearized version of the Bradford assay (Hayama et al. 2017). When making calibration curves, different quantities of known standard are used in order to infer the quantities of the substance of interest in a sample. We explored the effect of serial dilutions in the activity of the variants, expecting a proportional drop in activity after each dilution. We do *did* not observe a strong lack of linearity. However, we observed lower enzymatic activity for the NL3F10H variant compared to the MBP-NL3F10H in the tested conditions (Figure 2.A). This difference could be explained by a stabilisation or folding enhancement effect by MBP, helping to recover more active NL. The MBP tag adds the advantage for possible secondary purification step, using an Amylose column which can be bound by MBP, however we did not test this scenario.

**Figure 2.**
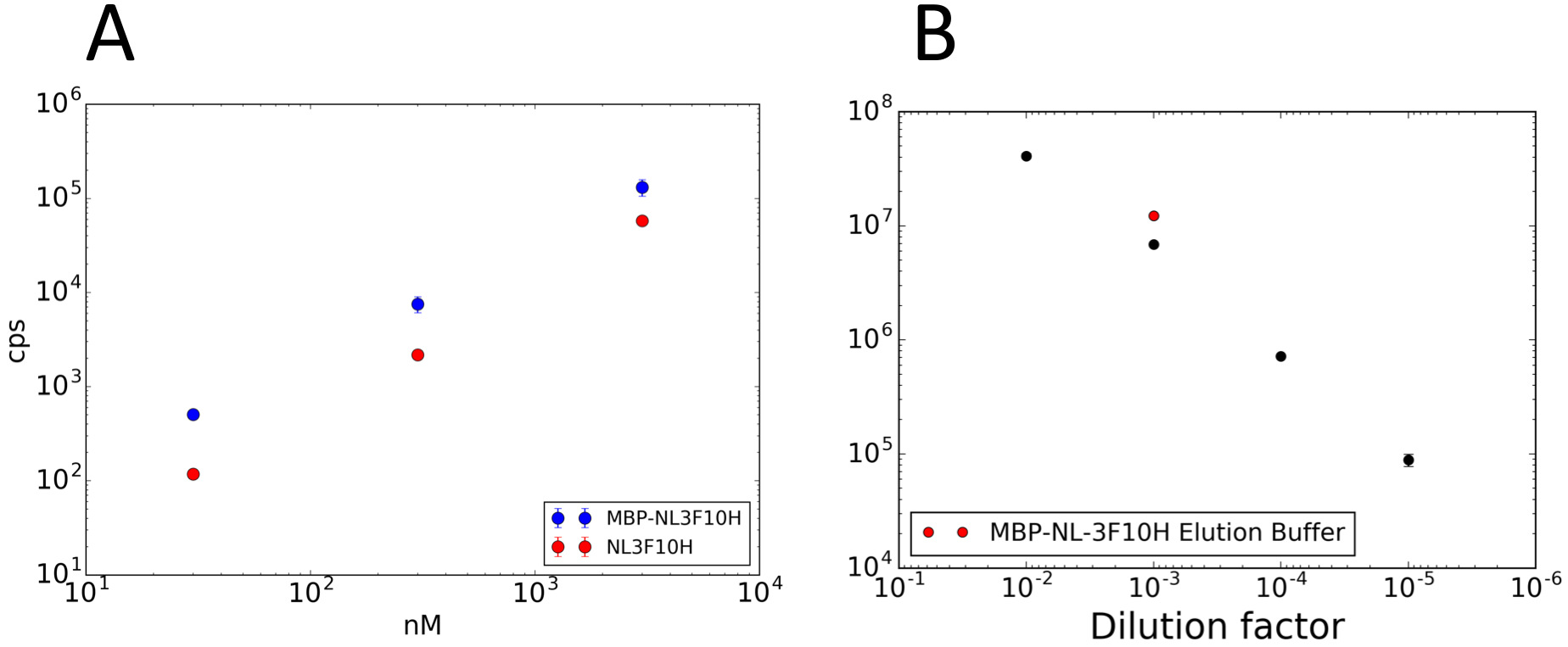
Behaviour of NanoLUC activity in dilutions. Purified NanoLUC versions were quantified by linearized Bradford assay and adjusted to 10mM. The reactions were incubated for 10 min at 21C and signal quantified in a Tristar plate reader with 1.5s signal integration time. The purification gels are presented in Figure 1. Error bars: S.E.M. counts per second (cps). A) panel dilutions in Elution buffer adjusted to the same molarity. B) MBP-NL-3F10H diluted in in Col-0 plant extract adjusted to 0.4 gFW/ml in Buffer BII. Red dot 1:1000 dilution in elution buffer (EB).

Then we explored whether plant extracts could inhibit MBP-NL3F10H activity by making serial dilutions of MBP-NL3F10H in a crude whole plant extract of 21-day-old Col-0 rosettes. The activity of NanoLUC is in this extracts that a 1×10^−3^ dilution was required for avoiding saturation problems with the plate reader (Figure 2.B). The results show that the signal decreases consistently by an order of magnitude after each dilution in plant extract. NanoLUC has higher activity in dilutions with Elution Buffer than with dilutions of plant extracts. This experiment highlights the importance of using *spike-in* extracts for building calibration curves that will be eventually used for inferring protein concentration in a sample to analyse. Working with an enzyme as a calibrator can be problematic if the activity is very unstable. If the enzyme decays very fast in pure preparation this will results in an overestimation of the tagged protein in freshly prepared plant extracts. Therefore, we did a simple experiment to test NanoLUC stability and derive a possible exponential decay model for the enzyme. This model can then be used for correcting quantifications because the enzyme is used within four days rather than immediately after purification. The results show that MBP-NL3F10H retains more than 95% of activity before three days and more than 80% after one week at 4°C. A linear regression on the log-transformed time series resulted in a half-life of 37.2 days (Figure 3). These results show that MBP-NL3F10H is a stable enzyme which can be used for generating calibration curves in plant extracts for absolute quantification. However, it is possible that there is room for improvement where very active preparation can be stored for very long periods of time without strong loss of enzyme activity.

**Figure 3.**
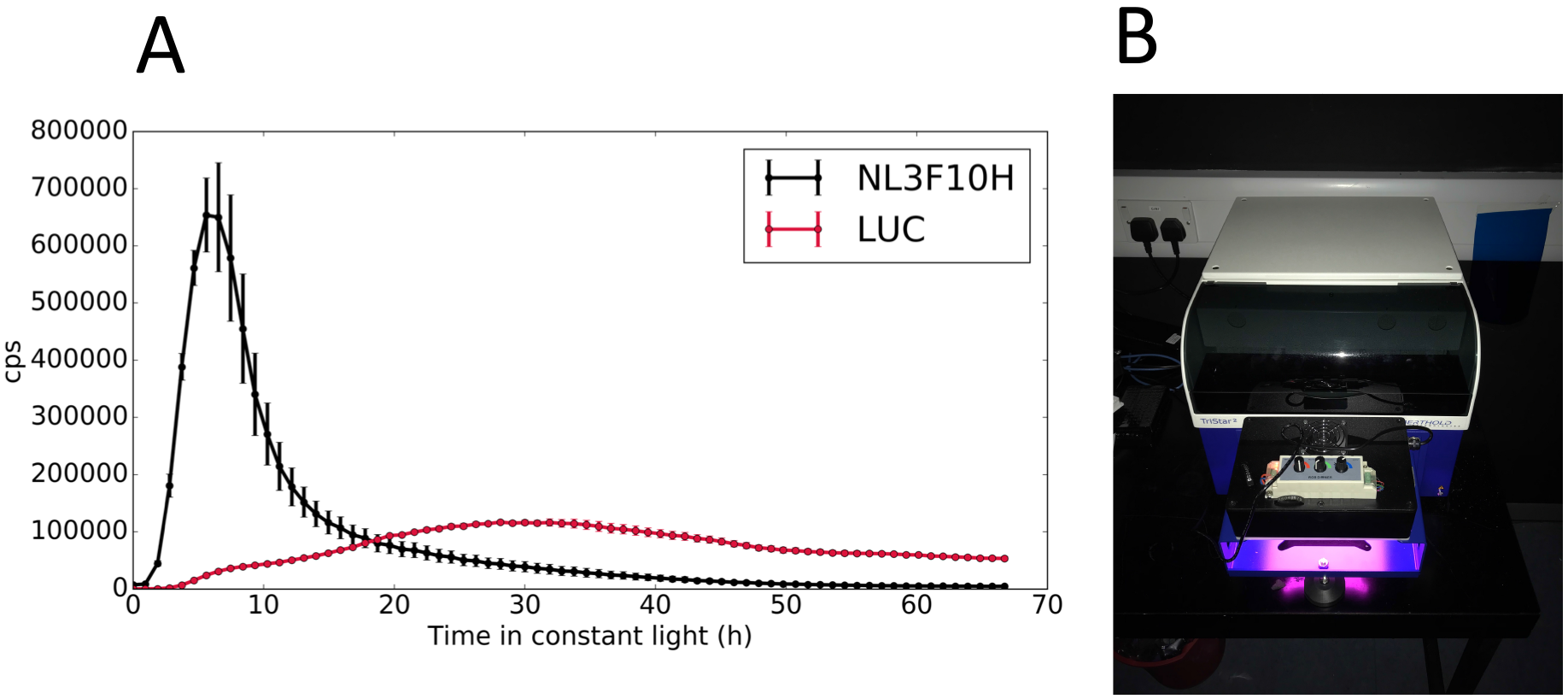
Activity of NanoLUC3F10H in protoplasts, compared to firefly luciferase (LUC). Protoplasts were isolated and transformed with equimolar amount of plasmid by the method of Hansen et al 2015. Then followed for three days in constant light at 21C 50 μmol/cm^2^s^2^ monochromatic blue and red LEDs in a 96-well black plate in a Tristar Berthold plate reader. Furimazine used as substrate for NL3F10H and luciferin for LUC. Mean of three technical replicates error bars show S.E.M.

### Extending pGWB with a versatile NanoLUC tag

The pGWB vector series is a good choice for the performing protein functional studies in plants. As it is a well stabilised vector series, we were able to focus on NanoLUC rather than in troubleshooting vector problems when agrobacterium mediated plant transformation (Nakamura et al. 2010; Nakagawa et al. 2007). An important characteristic of this collection is that they present a tail to tail construct:marker design, which might help to mitigate unintended effects on the expression of the synthetic construct by the selection marker. Second, the promoter on the selection marker is a NOPALINE SYNTHASE (NOS) promoter rather than a CaMV35S promoter. It has been shown that the CaMV35S promoter can result in secondary effects on the expression of nearby genes, possibly affecting the expression of the construct of interest (Yoo et al. 2005). Third, an extensive library of Gateway clones exists for plant systems therefore easing the use of NanoLUC without the need to re-clone this DNA parts. Finally, they present a good diversity of selection markers (Kanamycin, Hygromycin, BASTA and Tunicamycin). This is an important feature as many single and higher-order mutants carry different selection markers and complementation experiments on these mutants are crucial for testing the functionally of the fusions. We used pGWB601 as a base vector and introduced NL3F10H by Gibson Assembly. This resulted in the new pGWB601NL3F10H, with BASTA resistance as a selection marker for plant transformation. The resulting vector allows the generation of C-terminal translational fusions with NL3F10H by classical Gateway cloning. The new cassette was sub-cloned into pGWB401, pGWB501 and pGWB701, resulting in a collection of vectors with different plant selection markers (Figure 4).

**Figure 4.**
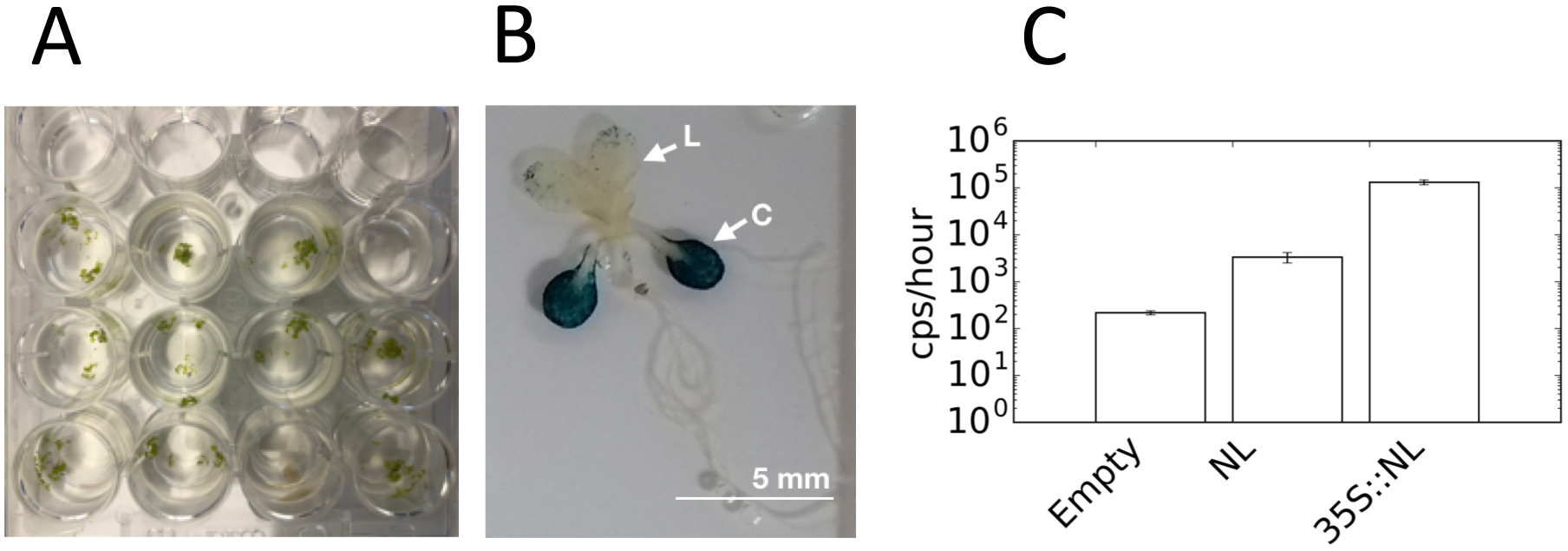
Testing NanoLUC using transient transformation in Arabidopsis seedlings using AGROBEST. A) Seedlings growing in ½ MES pH 5.4 0.5% sucrose are then transformed with Agrobacterium bearing Binary plasmids carrying constructs to be prototyped. B) Transformed seedlings using pCAMBIA1503.1 that contains 35S∷GUS, the GUS gene contains an intron that rules out GUS expression coming from Agrobacterium, Leaf (L), Cotyledon (C) C) Quantification of NanoLUC transformed seedlings luminescence was assessed by transferring seedlings to a 96 well flat white plate with 50 μl of 1:50 furimazine:0.01% Triton X-100 and tracked for 48 hours in constant light Error bars S.D. of time series for a single pool of 10 seedlings.

### Testing NanoLUC activity in Arabidopsis

In order to test the activity of NanoLUC in Arabidopsis, we cloned the CaMV35S (35S) promoter from the pCambia1305.1 vector into a pDONR221 vector for subsequent recombination into vectors pGWB601NL3F10 and pGWB635. We use the latter vectors using 35S:LUC+ constructs as positive control helping use to rule out problems with either the 35S sequence of transformation either in protoplast or plants. We performed a transient experiment with protoplast from 4-week old Col-0 plants, with megapreps of vectors containing 35S:NL3F10H and 35S:LUC constructs as described by (Hansen & van Ooijen 2016). We followed the signal for three days three days in constant light conditions at 21°C, in a Tristar plate reader (Figure 5.A). These results show that NL3F10H is translated into active enzyme in protoplasts. Interestingly the dynamics of the two constructs differ. NL3F10H presents a very sharp activation then decays. It is possible that the strong over-expression of NL3F10H results in substrate depletion compared to the LUC, however we did not explore this hypothesis. The NanoLUC signal can be detected earlier than LUC, possibly because the former is brighter. In addition to protoplasts, we also tested NanoLUC using the AGROBEST method. It allows transient transformation of *Arabidopsis* seedlings in an agrobacterium dependent manner (Wu et al. 2014). Four days after germination Col-0 efr-1 seedlings were co-incubated with Agrobacterium ABI strains, either carrying a vector without 35S promoter of a 35S:NL3F10H constructs. Also, Agrobacterium not carrying plasmids was used as negative control. The efr-1 mutation sensitizes Arabidopsis which otherwise is immune to *Agrobacterium* transformation through leaf infiltration. We observed strong bioluminescence in the 35S:NLF310H signal compared to the controls. However, we also detected signal in the pGWB601:NL3F10H vector without promoter, an order of magnitude lower though. This data suggested that the new vectors are competent for agro-mediated transformation in *Arabidopsis*.

**Figure 5.**
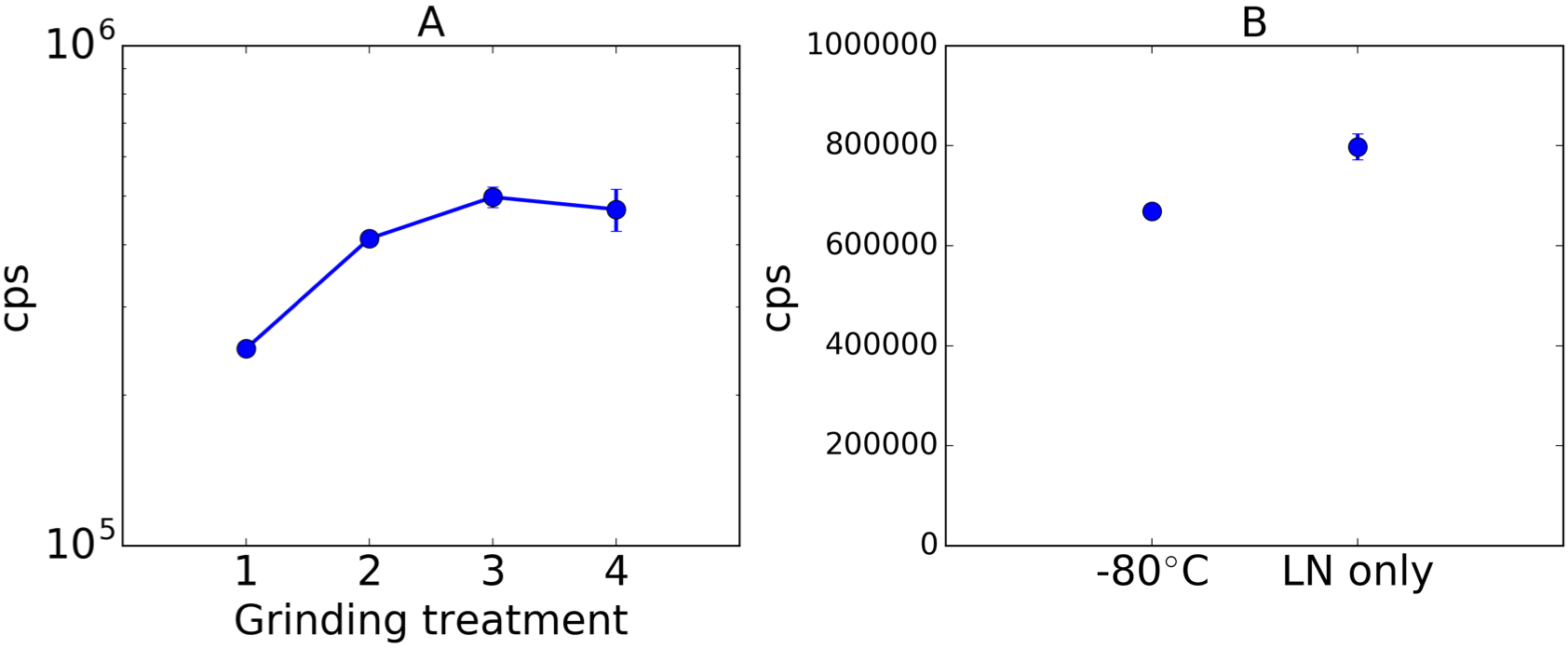
Extraction of NL3F10H from tissue of stable transgenic lines. A) Comparison of extraction method, 1) manual grinding, 2) treated once with liquid nitrogen and grinded in a Tissuelyser (Qiagen). 3) Second round of grinding in tissue lyser. 4) Third round of grinding of tissue lyser. B) Activity decays after storing tissue at −80°C for two days compared to liquid nitrogen (LN). 10 μl of plant extract mixed with 190 μl of BSII buffer then loaded in a 96-well black plate. Circles represent the mean three technical replicated with error bars representing SEM.

**Figure 6.**
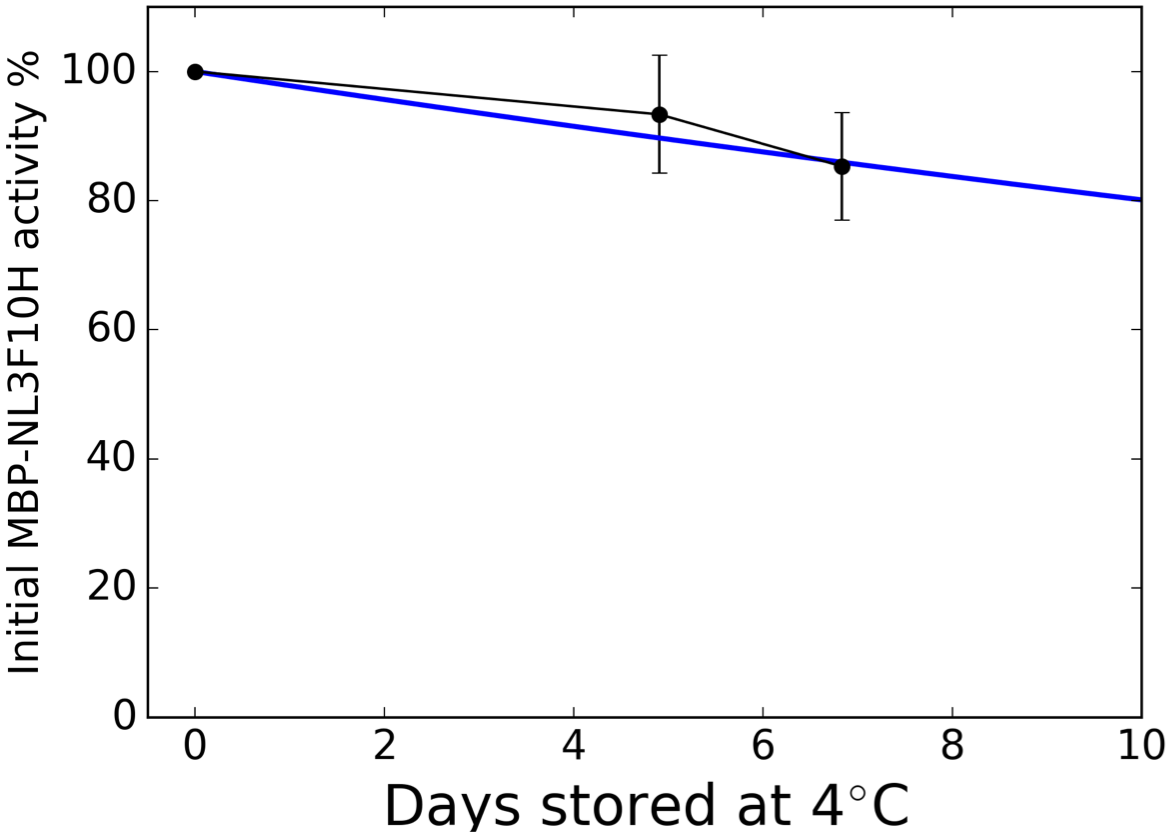
Decay of MBP-NL3F10H stored at 4 C in 250 mM NaCl 50mM NaH_2_PO_4_ pH 8.0 (NaOH adjusted). The purified enzyme was diluted 1:1000, then activity determined by mixing 20 μl of diluted enzyme with 80 μl of BI buffer and then mixing with 100 μl of 1:50 Furimazine:NanoGlow buffer in a 96-well flat black plate (Lumitrac, Greiner). Samples were incubated for 10 min at 21<u>°</u>C and measured with Tristar plate. Circles, mean values for three technical replicates, errors bars S.E.M. Solid blue line, fitted exponential decay models with results in a t_1/2_~37 days.

#### Overexpressing lines for optimisation of NanoLUC quantification in plant extracts

We transformed Col-0 plants by floral dipping to generate a collection of stable transgenic lines using the pGWB601:35S:NL3F10H vector. Resulting in a collection of lines that can be used as positive controls of NanoLUC activity. Also, the lines can also be used as negative control if NanoLUC-3Flag-10His is used as epitope tag for example in Chip-seq or Co-IP. Using BASTA resistance as marker we performed segregation analysis selecting lines that present single insertions judged by Mendelian ratios (3:1). Then, we chose randomly a line from the collection and evaluated the effect on NanoLUC activity when preparing plant extracts. First by manual grinding or by several rounds of tissue disruption mechanically (Figure 5.A.1). We observed that manual grinding with a pestle is less efficient compared to one or more round of mechanical extraction. In particular after the second round of grinding using Tissue Lyser (Qiagen) activity extraction saturates (Figure 5.A.3). This grinding experiments were performed with plants only treated with liquid nitrogen. However, in many experiments saving the samples for later experiments at −80 °C is a usual step (e.g. RNA isolation). Therefore, we tested this situation by flash freezing 2-weeks old seedlings expressing NanoLUC then storing them at −80 °C overnight. We observed decay in activity after storage vs processing them right after flash freezing them (Figure 5.B). This suggests that the samples should be processed on the same day or stored in liquid nitrogen if a time series is generated. However more replications should be conducted to in order to properly optimised storage conditions. For the moment in future work I will be conservative and process the samples without storing them at −80°C for later work.

### BOA-NL and BOA-LUC present different dynamics

Our experimental results show that NanoLUC is active in plants when used in several contexts. One of the reasons we are using NanoLUC is its reported brightness *in-vitro*, which has been reported to be 1000x brighter than LUC. The other strong reason to use NanoLUC is its stability which might in principle facilitate faithful tracking of protein levels *in vivo*. We decided to study the dynamics of a new component incorporated in the latest mathematical model F2014 is *BROTHER OF LUX ARRHYTHMO (BOA)*, also known as NOX. We cloned the genomic region of BOAp:BOA and created a translation fusion with NL3F10H using the pGWB601NL3F10 vector. Then transformed Col-0 CCA1p:LUC plants in order to see the effect of BOA-NL3F10 on the clock. BOA has been suggested to be an activator of CCA1 transcription which should be detectable in this background. We also created a BOAp:BOA-LUC construct and transformed Col-0 genotype to compare NL and LUC.

We performed a plate reader experiment with two independent lines for each construct (Figure 8). We used Col-0 CCA1p:LUC as a positive control for phase and period. BOAp:BOA-LUC reflects the expression pattern previously reported for the transcript by qRT-PCR (Figure 8.B). The signal reaches its highest levels late at night then drops to the lowest levels at mid-day in entrainment conditions. We observe that BOA presents rhythmic circadian oscillations, CCA1p:LUC and BOA-LUC have antiphase relationship. It is worth noting that in the two BOA lines selected, the signal from BOA is lower than what *that of* CCA1p:LUC. This might indicate that BOA is expressed at low levels. Dai et al (2011) reported that BOA can *be* detected just above the limit of detection using antibodies against the native protein. This is consistent with a low signal of LUC, however this may be specific to the two lines selected. In the same experiment we also monitored the levels of BOAp:BOA-NL3F10H finding that BOA-NL3F10H protein does not oscillate with high amplitude. This comes as a surprise because the F2014 model predicts oscillatory behaviour (Fogelmark & Troein 2014). However we observe a correlation between NanoLUC signal and period of CCA1p:LUC which indicates that the fusion is genetically active and consistent with observations that overexpressing BOA results clock period lengthening (Figure 8.C). In principle the only difference between BOA-NL and BOA-LUC is the reporter sequence. However, this data might indicate that differences in stability of NL and LUC explain the waveform differences. We created a simple ODE model, to explore how different parameters might explain the waveform.

**Figure 7.**
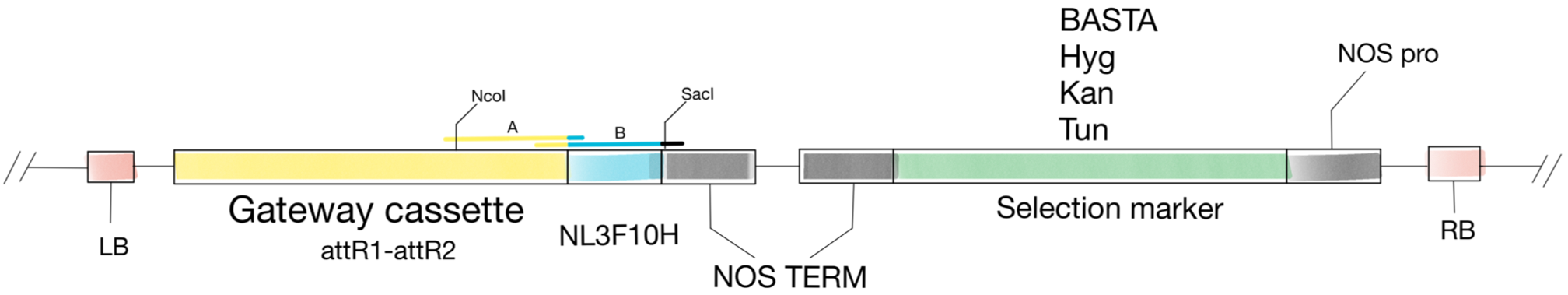
Structure of pGWB vectors extended for NL3F10H translational fusions. The Gateway NL3F10H cassette was generated by Gibson Assembly of PCR products A and B, designed with overlapping regions between them and pGWB601 vector. The vector was digested with Nco/SacI restriction enzymes to insert the new cassette. T-DNA Left Border (LB) and right border (RB). NOPALINE SYNTHASE promoter and terminator (NOS pro, NOS Term). Gateway cassettes of type attR1-attR2 can be used as destination in a LR Gateway cloning reaction. The pGWB vectors have a common Scp^r^ marker for selection in bacteria, with a pPZP vector backbone. The variants used here were pGW401 (Kan^r^), pGWB501 (Hyg^r^), pGWB601 (BASTA^r^) and pGWB701 (Tun^r^). NL3F10H was introduced in each of them by NcoI/SacI sub-cloning.

**Figure 8.**
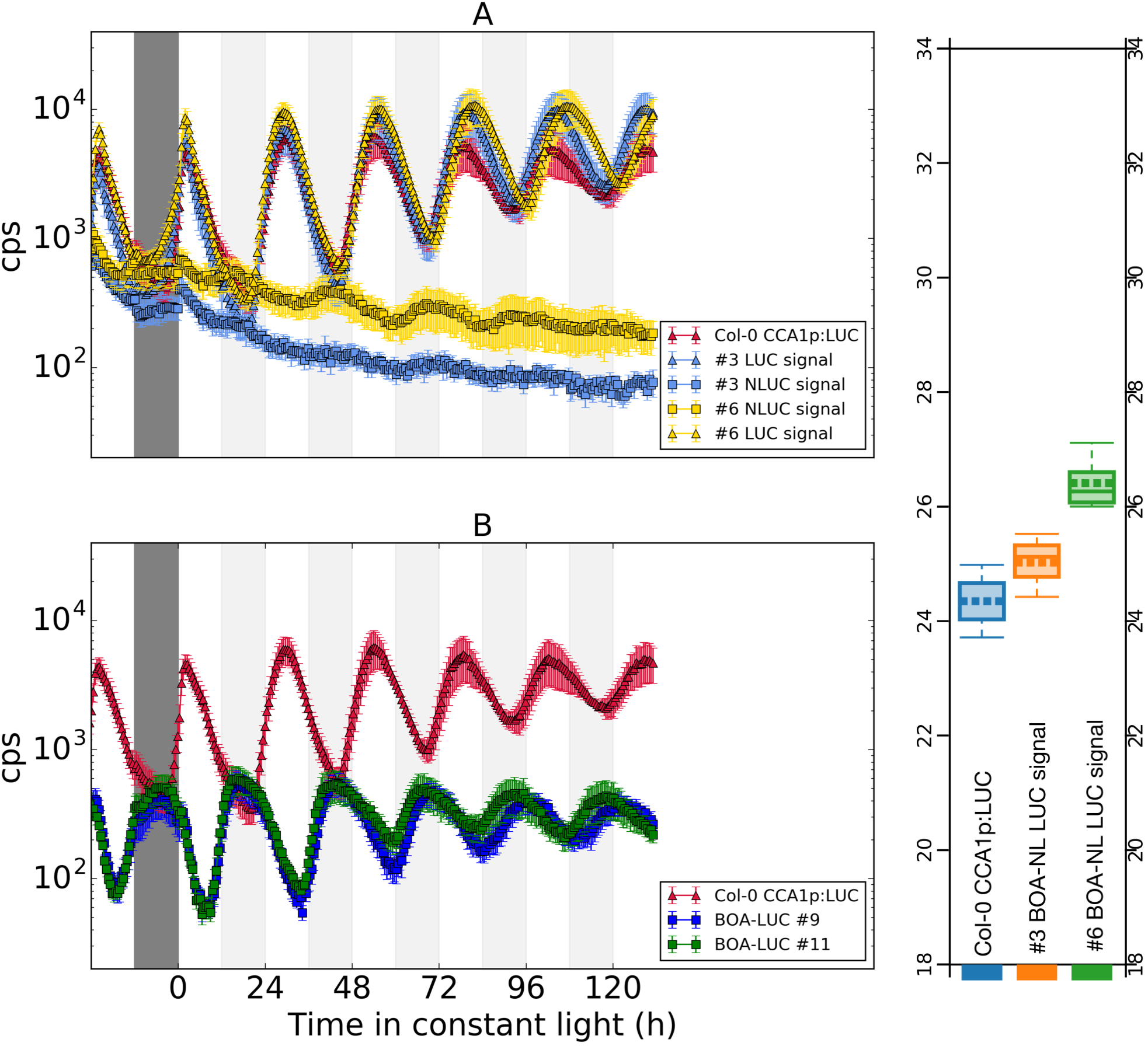
Dynamics BOA protein using NanoLUC and LUC reporters. A) Dynamics in Col-0 CCA1p:LUC BOAp:BOA-NL3F10H using two independent T3 lines, The signal of CCA1p:LUC (triangle) and BOA-NL3F10H (squares) were measured in different wells, the control is shown in red. B) Dynamics of BOAp:BOA-LUC in Col-0. Seedlings were grown for 7 days at 140 μmol/cm^2^s^2^ white light 12L:12D 21°C. Then transferred for 2 days into 50 μmol/cm^2^s^2^ monochromatic blue and red LEDs, before quantification started using a Tristar plate reader every 30 min with 1.5s of signal integration. The traces are means of 4 replicates assays error bars represent SEM. C) BOA-NL impacts the period of CCA1p:LUC. Y-axis period estimates for time-series in A. Period determined in Biodare2 using FFT-NLS method.

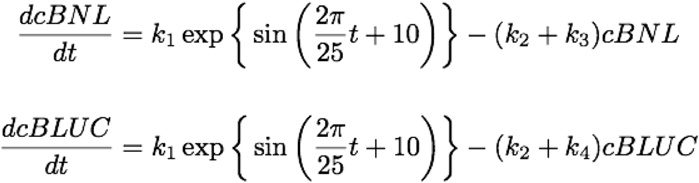

Where *cBNL* and *cBLUC* are state variables for BOA-NL3F10H and BOA-LUC. *k*_*1*_ is the translation rate that is identical between the two constructs, *k*_*2*_ is the degradation rate BOA protein, *k*_*3*_ the decay rate of NL3F10H activity and *k*_*4*_ the decay rate of LUC activity. The decay rate of NL3F10H was fixed to the one determined at 4°C in this work, however the decay rate was adjusted for temperature assuming a Q_10_=2.5 and a temperature of 21 given that the plate reader experiment was conducted at that temperature. k_1_, k_2_ and k_4_ were adjusted manually for matching the experimental data (Figure 9). The model can capture the waveform, furthermore we observed a decay constant for firefly LUC in a realistic value (Table 1).

**Figure 9.**
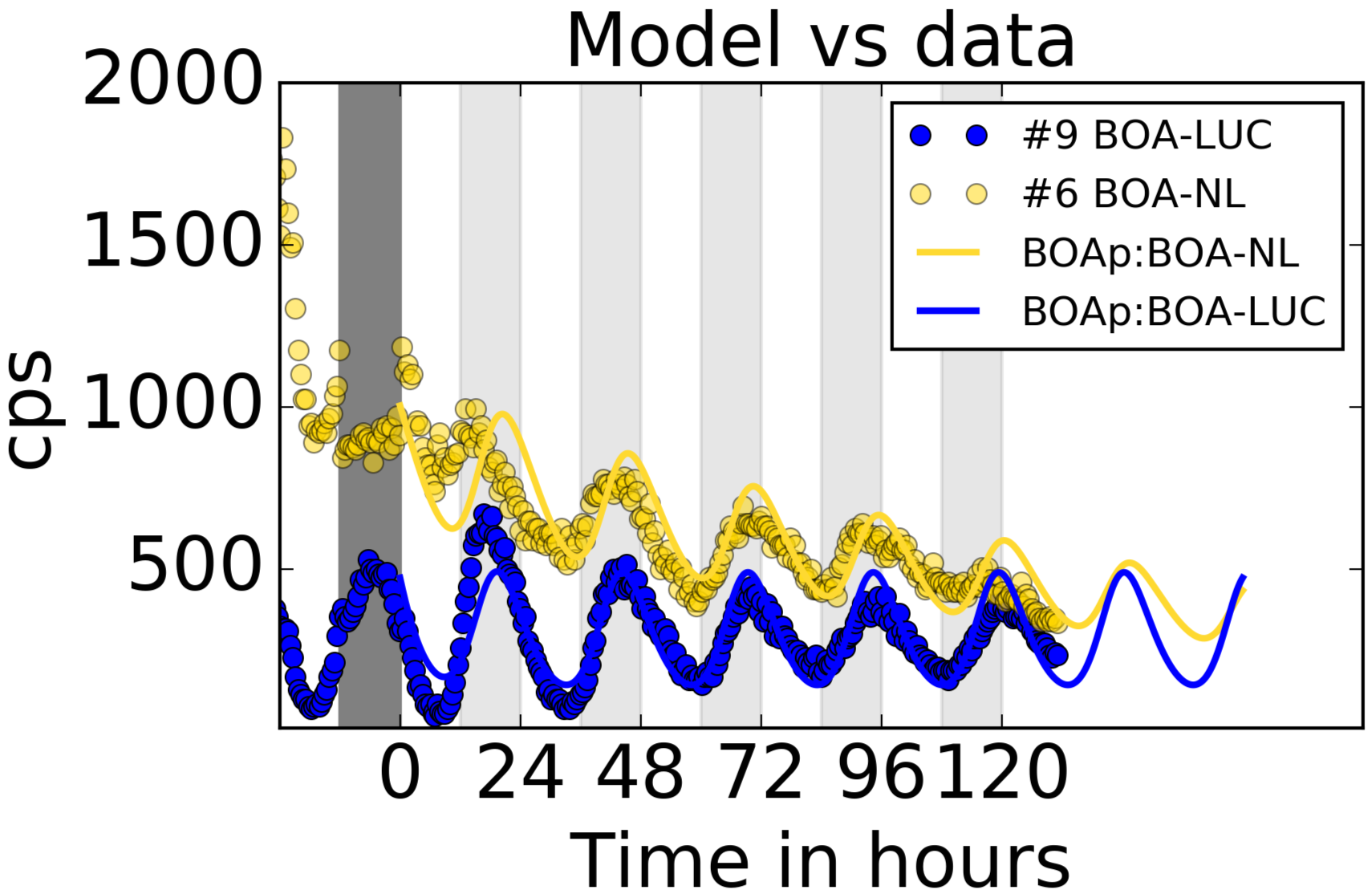
Comparing model vs luminescence data. The brightest traces of BOA-NL and LUC for the indicated lines are presented (circles). The model from Table 1 is in solid lines.

**Table 1.** Model parameters for the simple model that assumes only differences in decay constants between reporters *K*_*3*_ (NanoLUC) and *K*_*4*_ (LUC). Fitting the parameters show that is possible to replicate the signal. NanoLUC decay rate was obtained by scaling the experimentally determined rate assuming a Q10=2.5 as plate reader experiments were conducted at 21^°^C. The trend could possibly be explained by decay in the Furimazine concentration, for which a exponential decay was introduced introduced.

| PARAMETER | VALUE | UNITS | SOURCE | PROCESS |
| --- | --- | --- | --- | --- |
| K1 | 50 | cps/h | Fitted | Translation |
| K2 | 0.08 | 1/h | Fitted | BOA decay |
| K3 | 0.0121 | 1/h | Experimental | NanoLUC decay |
| K4 | 0.15 | 1/h | Fitted | LUC decay |
| A | 0.004 | 1/h | Fitted | Furimazine decay |

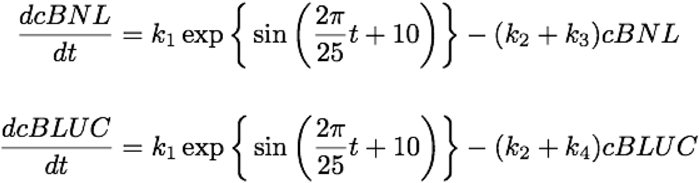

## Discussion

The use of tagged lines with general epitopes e.g. GFP, FLAG 10xHis has been a powerful approach to study the protein component of the *Arabidopsis* circadian oscillator. Thanks to these lines, several aspects of clock protein regulation have been investigated, including localisation, binding sites in the genome by ChIP, and protein-protein interactions by Co-IP or, more recently, affinity purification mass spectrometry (e.g. the case of ELF3 and GI. (Huang et al. 2016; Krahmer et al. 2017)). We decided to explore the use of NanoLUC as possible new reporter for circadian biology in plants. We showed that it is possible to work with NanoLUC in infrastructure developed for tracking LUC bioluminescence. We hope this work provides the community with a set of initial guidelines for working with NanoLUC, at least in plants. We report a purification protocol that can be used for generating calibration curves using NanoLUC. Results also demonstrate that the protein is active in either in transient expression systems and stable transgenics. We developed a collection of 35S:NL3F10H that can be used as positive controls for NanoLUC signal or for negative controls if NanoLUC-3F10H is used as an epitope.

We tested NanoLUC in a real case scenario by generating new protein data for BOA. Surprisingly we observed a discrepancy between NanoLUC and LUC. Both constructs were generated from the same pDONR221:BOApBOA-no-stop vector therefore the only difference is the reporter tag. LUC presented clear oscillations in light emission, although these are low compared to CCA1p:LUC. Only one BOA NanoLUC line presented subtle oscillations. However, in BOA NanoLUC lines presented period changes compared to the Col-0 CCA1p:LUC+ which suggest that BOA-NL3F10H is active in these lines. One possibility for the difference in behaviour is a difference in the stability of BOA-LUC and BOA-NL3F10H. In particular the instability of LUC has been exploited as a way to track transcriptional rhythms. We wanted to exploit NanoLUC because its stability might help to provide a representation of total protein levels. A simple mathematical model was the only difference between LUC and NanoLUC is the decay rate can explain the differences between the two reporter designs. Using the experimentally determined NanoLUC decay rate we inferred a half-life of 8.6 h for BOA and of for LUC of 4.6 h. In mammalian cells LUC has a 2-hour half-life at 37°C, correcting the half-life by using the Q10 of 2.5 results in 5 hours, therefore it seems that we are obtaining a reasonable estimate for LUC half-life, consistent with BOA being a stable protein. However, this comes as a consequence of assuming equal brightness between NanoLUC and LUC *in-planta*. The lower NanoLUC activity observed *in-planta* do not seem to be a consequence of strong inhibitors in cytosol as we did not observe a strong decay in signal when mixing pure NanoLUC with plant extracts of Col-0. This indicates that the permeability of furimazine is the current bottle neck for increasing sensitivity *in-planta* protein tracking. This will become relevant if researchers want to use NanoLUC for tracking protein levels using optical systems, though proper tracking of NanoLUC fusions *in-planta* using sensitive photomultipliers still a viable option. Overall our results indicate that NanoLUC is compatible with stablished methodologies for LUC in *Arabidopsis* and we expect to see increased use of this luciferase for gathering high-fidelity quantitative protein data in plants.

## Declarations

### Author’s Contributions

U-UG and AJ-M planed experiments, analysed data, U-UG executed experiments and wrote the manuscript.

## Acknowledgements

We thank Eilidh Smith assessing manuscript readability

## Competing Interests

The authors declare that they do not have competing interests

## Availability of data and materials

NanoLUC is property of Promega use of any material containing NanoLUC requires license acquisition

## Funding

The research was funded by Consejo Nacional de Ciencia y Technologia (CONACY), Mexico, The School of Biological Sciences the University of Edinburgh PhD scholarship

## References

Bognár, L. et al., 1999. The circadian clock controls the expression pattern of the circadian input photoreceptor, phytochrome B. Proceedings of the National Academy of Sciences of the United States of America, 96(25), pp.14652–14657.

Chow, B.Y. et al., 2012. ELF3 recruitment to the PRR9 promoter requires other Evening Complex members in the Arabidopsis circadian clock. Plant signaling \& behavior, 7(2), pp.170–173.

Corellou, F. et al., 2009. Clocks in the green lineage: comparative functional analysis of the circadian architecture of the picoeukaryote ostreococcus. The Plant cell, 21(11), pp.3436–3449.

Dai, S. et al., 2011. BROTHER OF LUX ARRHYTHMO is a component of the Arabidopsis circadian clock. The Plant cell, 23(3), pp.961–972.

Feeney, K.A. et al., 2016. Daily magnesium fluxes regulate cellular timekeeping and energy balance. Nature, 532(7599), pp.375–379.

Fogelmark, K. & Troein, C., 2014. Rethinking transcriptional activation in the Arabidopsis circadian clock. PLoS computational biology, 10(7), p.e1003705.

Hall, M.P. et al., 2012. Engineered luciferase reporter from a deep sea shrimp utilizing a novel imidazopyrazinone substrate. ACS chemical biology, 7(11), pp.1848–1857.

Hansen, L.L. & van Ooijen, G., 2016. Rapid Analysis of Circadian Phenotypes in Arabidopsis Protoplasts Transfected with a Luminescent Clock Reporter. JoVE (Journal of Visualized Experiments), (115), pp.e54586–e54586.

Hayama, R. et al., 2017. PSEUDO RESPONSE REGULATORs stabilize CONSTANS protein to promote flowering in response to day length. The EMBO journal, 36(7), pp.904–918.

Huang, H. et al., 2016. Identification of Evening Complex Associated Proteins in Arabidopsis by Affinity Purification and Mass Spectrometry. Molecular & cellular proteomics : MCP, 15(1), pp.201–217.

Jinek, M. et al., 2012. A programmable dual-RNA-guided DNA endonuclease in adaptive bacterial immunity. Science (New York, N.Y.), 337(6096), pp.816–821.

Krahmer, J. et al., 2017. Time-resolved Interaction Proteomics of the Putative Scaffold Protein GIGANTEA in Arabidopsis thaliana. bioRxiv, p.162271.

Millar, A. et al., 1995. Circadian clock mutants in Arabidopsis identified by luciferase imaging. Science (New York, N.Y.), 267(5201), pp.1161-1163.

Millar, A., Short, S., Chua, N., et al., 1992. A novel circadian phenotype based on firefly luciferase expression in transgenic plants. The Plant cell, 4(9), pp.1075-1087.

Millar, A.J., Short, S.R., Hiratsuka, K., et al., 1992. Firefly luciferase as a reporter of regulated gene expression in higher plants. Plant Molecular Biology Reporter, 10(4), pp.324-337.

Nakagawa, T. et al., 2007. Improved Gateway binary vectors: high-performance vectors for creation of fusion constructs in transgenic analysis of plants. Bioscience, biotechnology, and biochemistry, 71(8), pp.2095-2100.

Nakamura, S. et al., 2010. Gateway binary vectors with the bialaphos resistance gene, bar, as a selection marker for plant transformation. Bioscience, biotechnology, and biochemistry, 74(6), pp.1315-1319.

Puigbò, P., Bravo, I.G. & Garcia-Vallvé, S., 2008. E-CAI: a novel server to estimate an expected value of Codon Adaptation Index (eCAI). BMC bioinformatics, 9(1), p.65.

Shen, J. et al., 2013. Organelle pH in the Arabidopsis endomembrane system. Molecular plant, 6(5), pp. 1419-1437.

Suter, D.M. et al., 2011. Mammalian genes are transcribed with widely different bursting kinetics. Science (New York, N.Y.), 332(6028), pp.472-474.

Wu, H.-Y. et al., 2014. AGROBEST: an efficient Agrobacterium-mediated transient expression method for versatile gene function analyses in Arabidopsis seedlings. Plant methods, 10(1), p.19.

Yoo, S.Y. et al., 2005. The 35S promoter used in a selectable marker gene of a plant transformation vector affects the expression of the transgene. Planta, 221(4), pp.523-530.

